# Application of benchmark analysis for mixed contaminant exposures: Mutual adjustment of two perfluoroalkylate substances associated with immunotoxicity

**DOI:** 10.1101/198564

**Authors:** Esben Budtz-Jøergensen, Philippe Grandjean,

## Abstract

**Background:** Developmental exposure to perfluorinated substances is associated with deficient IgG antibody responses to childhood vaccines as an indication of depressed immune system functions. As this outcome may represent a critical effect of these substances, calculation of benchmark dose (BMD) results would be useful for standards setting to protect exposed populations against adverse effects. However, in the mixed exposure setting of most epidemiological evidence, the two major and inter-related substances associated with this adverse effect have shown similar benchmark results that raise concerns about possible confounding.

**Methods:** With the aim of better characterizing the immunotoxicity impact of the two major perfluorinated substances in the mixed exposures, we carried out BMD calculations on prospective data from two prospective birth cohort studies from the Faroe Islands with a total of 1,146 children. Exposure data included serum concentrations of perfluorooctane sulfonate and perfluorooctanoate at birth and at age 5 years and, as outcome parameters, the serum concentrations of specific IgG antibodies against tetanus and diphtheria at ages 5 and 7. We calculated the BMDs and their lower confidence bounds (BMDLs) and included mutual adjustment for the two compounds.

**Results:** The BMDLs for the two immunotoxicants were of similar magnitude before and after adjustment. Both substances showed lower results for a logarithmic dose-response model, which also provided a slightly better fit than a linear dose model for both antibodies. We also used a broken curve shape that allowed a different slope below the median exposure. Postnatal exposure as represented by the age 5 serum concentration, showed a stronger association with the antibody outcomes than the prenatal exposure. Due to the correlation between the two immunotoxicants, the mutual adjustment resulted in elevated BMD results and *p* values. However, the BMDL values were virtually unchanged.

**Conclusions:** Adjustment for co-exposure to another immunotoxicant increased the variance and the BMD values, but affected the BMDL values only to a negligible extent. These calculations are in accordance with an interpretation that, when two toxicants appear to affect an outcome to an almost equal degree and none of them is known to be solely responsible, the exposures should both be considered responsible and attract equal regulatory attention until further evidence shows otherwise.

## Introduction

Perfluorinated alkylate substances (PFASs) have been in use for over 60 years in a wide array of applications [1], but it was not until the beginning of this millennium that academic research began to focus on their environmental fate, human exposures, and possible adverse effects [2]. Of note, an internal industry-commissioned study from 1978 revealed immunotoxicity in monkeys exposed to perfluorooctane sulfonate)PFOS) [3]. When these results were released to the U.S. Environmental Protection Agency (document AR 226-0139) many years later, they helped inspire laboratory studies to elucidate PFAS immunotoxicity [4-6].

By now, PFAS immunotoxicity has been demonstrated in rodent models, avian models, reptilian models, and mammalian and non-mammalian wildlife [7]. For example, in a commonly used mouse model, effects of perfluorooctanoate (PFOA) include decreased spleen and thymus weights, decreased thymocyte and splenocyte counts, decreased immunoglobulin response, and changes in specific populations of lymphocytes in the spleen and thymus. Reduced survival after influenza infection has been reported as an apparent effect of PFOS exposure in mice [8]. In one model, the lowest observed effect level for males corresponded to an average serum-PFOS concentration of 92 ng/mL, although 7-fold higher in females [5]. The former is similar to the highest concentrations found in serum from humans with background exposures [9].

The experimental findings triggered epidemiological studies to assess sensitive markers of immune functions. As recommended by an international group of immunotoxicity scientists, concentrations of specific antibodies against childhood immunizations were deemed to be a both feasible and appropriate outcome parameter, as children receive the routine immunizations with the exact same doses and at approximately the same age [10]. This approach would also take into account developmental vulnerability, as the first routine immunizations are usually given in early infancy when the evolving adaptive immune system is sufficiently capable of responding to antigen challenges.

Population-based serum-PFAS analyses show that PFOS and PFOA are detectable in virtually all Americans [9], with children often showing higher serum concentrations than adults [11]. Paired samples of maternal serum and cord serum show that PFASs are transferred through the human placenta [12, 13], and these substances also occur in human milk [14]. Thus, long-term breastfed infants may reach serum-PFAS concentrations that are several-fold higher than the mother’s [14, 15]. The focus on vaccination responses in children vaccinated in infancy therefore appears to be highly relevant to risk assessment.

We prospectively followed birth cohorts in the Faroe Islands and showed that developmental PFAS exposures constitute a major determinant for antibody concentrations directed against tetanus and diphtheria toxoids at ages 5 and 7 years, i.e., after three or four routine vaccinations [16, 17]. Our most recent findings suggest that elevated serum-PFAS concentrations in infancy are particularly associated with lower antibody concentrations at age 5 [17]. In contrast, at age 7 years, the postnatal PFAS accumulation seems to play a major role [16]. In agreement with this observation, antibody responses to vaccinations at adult age have also been reported to be lower at elevated serum-PFAS concentrations [18, 19].

When analyzing dose-dependent adverse effects, regulatory agencies often use benchmark dose (BMD) calculations to obtain standardized points of deviation for deriving safe exposure limits [20, 21]. For example, basic PFAS toxicity data obtained from animal models have been applied to calculate BMDLs and at first resulted in very high levels that corresponded to serum concentrations of 23 μg/mL and 35 μg/mL for PFOA and PFOS, respectively [22-24]. Even when taking into account large uncertainty factors, a calculated safe serum concentration would greatly exceed current levels in humans [9]. Given that immunotoxicity has been observed at current exposure levels [16], the BMDL results obtained from routine laboratory toxicity tests are clearly insufficient to serve as a basis for safety calculations, as has also been seen in regard to other pollutants [25]. Although substantial evidence on immunotoxicity in animal models is now available [7], BMD calculations have apparently not been reported.

So far, human data on PFAS toxicity have not been considered by regulatory agencies for determining exposure limits, e.g., because of the absence of a non-exposed control group [26]. In a recent evaluation, the National Toxicology Program (NTP) considered the epidemiological evidence for PFOA and PFOS immunotoxicity to be only moderate, given that all studies are observational and relate to mixed exposures [27]. As both PFOS and PFOA show adverse effects on the same target, published data do not allow a judgment if only one of them is the culprit. However, our recent calculations show that the effects of PFOS and PFOA appear to be independent, at least in part, although the statistical power was insufficient to allow an assessment of potential interactions between the two exposures [28]. *In vitro* data using human white blood cells support the immunotoxic potential for both PFOS and PFOA and that the adverse outcome pathways may differ [29].

On this background, the present report extends previous BMD results for PFAS-associated immunotoxicity [30] using an extended data base from two birth cohorts, with a wide range of background exposures both prenatally and postnatally, now including mutual adjustment for concomitant PFAS exposures. Because of the similarity to exposures in other populations [9], our results should be applicable beyond the Faroes. As before, the choice of dose-response models must take into account the absence both of a known curve shape and a null exposure control group. We therefore explore several options to elucidate the impact of extrapolations beyond the exposure interval observed [31].

## Methods

### Birth cohort data

Our studies rely on birth cohorts from the fishing community of the Faroe Islands, where PFAS exposures in part originate from marine food [32]. The oldest birth cohort in this study was recruited in 1997-2000 and consisted of 656 singleton births [16]. Prospective follow-up included 587 cohort members participating in one or both examinations at ages 5 and 7 years, of whom 460 participated in both, and complete serum analyses were obtained for 431. As an exposure indicator, we used the PFAS concentrations in the mother’s pregnancy serum and in the child’s serum obtained at age 5 years. The outcomes were the specific antibody concentrations against tetanus and diphtheria toxoids in serum at ages 5 and 7 years.

The younger birth cohort consisted of 490 children born during 2007-2009 [33], so far follow-up only through age 5 years. A maternal serum sample was collected shortly after childbirth to represent the child’s prenatal exposure, and serum-antibody concentrations were obtained at age 5 (before the booster) in 349 cohort members. As comparable methods were used, we were able to combine the results from the two cohorts regarding prenatal exposure and antibody results at age 5.

Among the PFASs measured in mothers and their children, PFOS and PFOA showed the highest concentrations [16, 33], similar to levels reported from the US [9]. Antibody concentrations showed substantial variability, many children with concentrations below the clinically protective level, despite having followed the recommended vaccination schedule.

Given the strong inverse associations between measured developmental exposures and specific antibody concentrations, we now report extended benchmark calculations first for the older cohort using the serum-PFAS concentration at age 5 as predictor of the antibody outcomes at age 7 years. These calculations correspond to our previous report [30], now supplemented by the results with mutual adjustments for PFOS and PFOA.

For the first time, we now also present the benchmark results for prenatal PFAS exposure in regard to the antibody concentration outcomes at age 5 years. These results are based on both cohorts, and again, the results for PFOS and PFOA are mutually adjusted.

The two sets of calculations represent prenatal and mid-childhood exposures as different windows of immune system vulnerability. As an advantage in this population, exposures to methylmercury and polychlorinated biphenyls were only weakly correlated with serum concentrations of the PFASs [16], and confounding by these other exposures could therefore be ignored.

### Benchmark calculations

The data were analyzed as continuous variables in SAS version 9.4 (SAS Institute, Inc. Cary, NC). Although a clinical cut-off level exists for antibody concentrations that represent long-term protection, this limit is somewhat arbitrary, and transformation of the continuous data to a dichotomous variable would result in a loss of information. Benchmark calculations were therefore based on regression models with antibody concentrations as dependent continuous variables while PFAS-concentrations were included as independent variables along with potential confounders. To achieve normally distributed residuals, antibody concentrations were log-transformed. Thus, the models were on the following form:

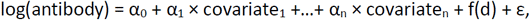

where d is the PFAS concentration (PFOS or PFOA) and f is the dose-response function satisfying f(0) = 0. We modeled the PFAS-effect using a linear-dose response function [f(d) = β × d], and a logarithmic model [f(d) = β × log(d+1)]. As the dose-response relationship at low doses may differ from the one at higher doses, we also used a piecewise linear model, which allowed for a difference in slopes below and above the median exposure level. The fit of the models was based on minus two times the log-maximum likelihood function (−2 log(L)), where a smaller value indicates a better fit. Benchmark calculations were first carried out for PFOS and PFOA separately, then adjusted models were developed where one of the serum concentration were included as the exposure of interest, while the other entered the model as a covariate.

Benchmark results were obtained for PFAS-concentrations measured at age 5 years and for prenatal exposures in the two sets of analyses, with antibody concentrations at ages 7 and 5 as the respective outcome variables. As previously observed [16], relevant covariates comprised sex, age and booster type at age 5 (the latter only for the age 7 outcomes). For the analyses of prenatal exposures, regression models included observations from both cohorts, and cohort was therefore treated as a covariate. We also allowed the effect of sex and age to differ in the two cohorts by including interaction terms.

The BMD is the dose which reduces the outcome by a certain percentage (the benchmark response, BMR) compared to the unexposed controls [20, 21]. Different BMR values have been used in the past, and lower BMR levels are known to result in decreased BMD and BMDL results, the latter affected by increased uncertainty [31]. A 10% BMR is usually applied for experimental toxicology data [20, 21], while in human studies a BMR of 5% is often chosen [20]. As a decreased antibody response to vaccinations must be regarded an important adverse effect, the lower BMR would seem appropriate. We calculated BMD results for both BMR values.

An advantage of a log-transformed response is that BMD can be estimated independently of the confounders as the dose where the dose-response function is equal to log (1-BMR), i.e., the BMD, will satisfy the equation f(BMD) = log (1-BMR). If the estimated dose-response is increasing (indicating a beneficial effect), the BMD is not defined. In the current study, such cases occurred when PFAS-exposures were mutually adjusted, and the BMD is then indicated with a 8.

The BMDL is defined as the lower one-sided 95%-confidence limit of the BMD [20, 34]. In our dose-response models with linear parameters, the derivation of closed-form expressions for the BMDL is straight forward [35]. Based on the estimated uncertainty in the parameter estimates, the lower confidence limit of the dose-effect function [f(d)] can be easily determined. The BMDL is given as the dose where this confidence limit is equal to log(1-BMR).

As a consequence of the relatively steep dose-response relationships, the BMDs were often lower than the minimum observed exposure. Consequently, some results depended on a part of the dose-response curve for which the data does not hold any information. As a sensitivity analysis, we therefore developed a low-dose threshold version of each of the dose-response models used. Each of these models was identical to the original dose-response model within the observed dose range, but with a flat dose-response slope below the lowest dose observed (Figure 1).

**Figure 1.**
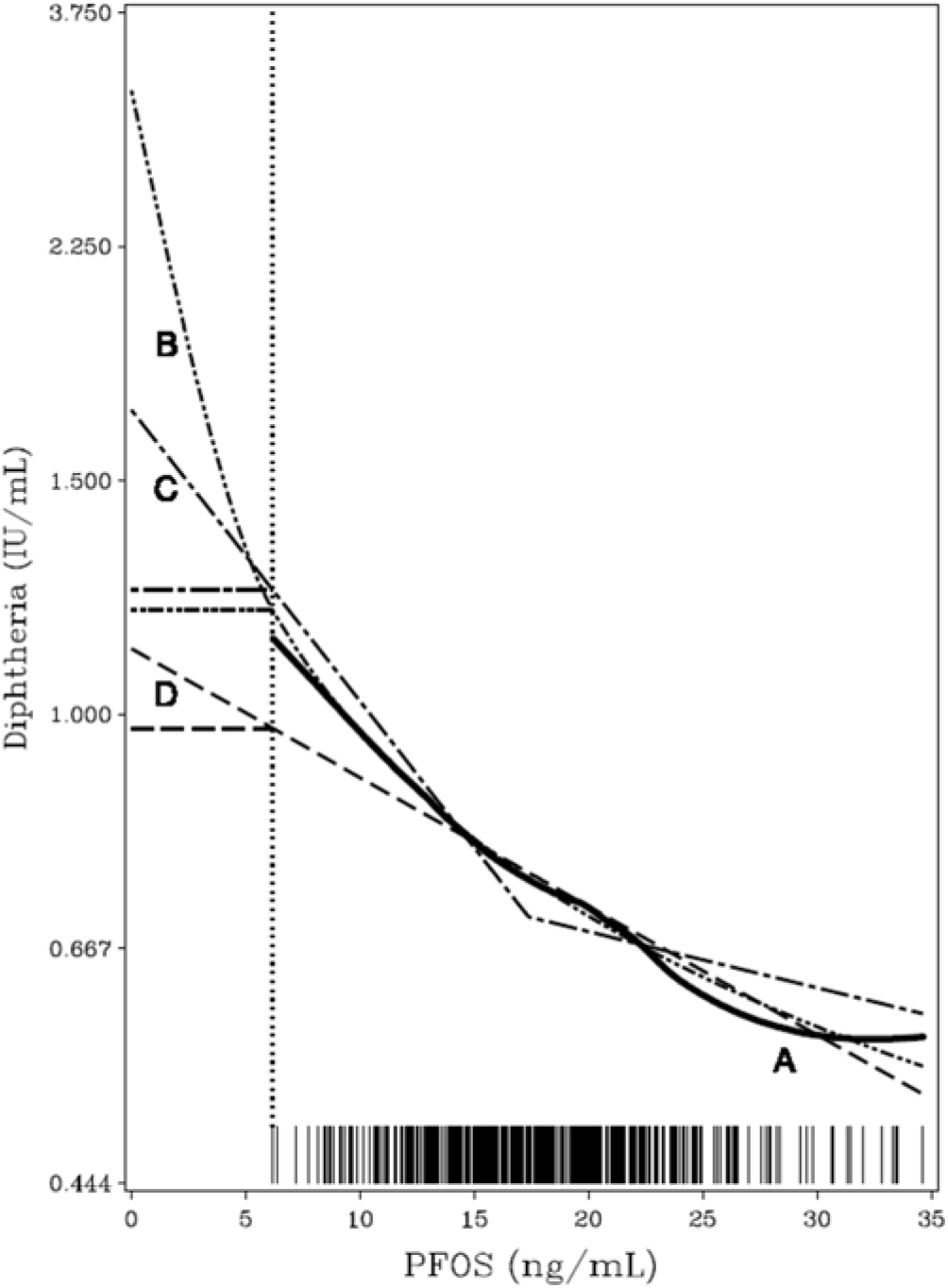
Estimated dose-response functions for the relationship between PFOS and Diphtheria. Curve A is estimated as a generalized additive model. Curve B is the log-function, C is piecewise linear and D is linear. The low-dose threshold models (see Table 3) assume a flat curve below the lowest observed dose indicated by the dotted vertical line, i.e., that a threshold exists at the lowest serum-PFC concentration observed [30].

## Results

Both cohorts were affected by a fairly small degree of attrition, but children who participated in one clinical examination, but not the other, or not at all, did not seem to differ in terms of exposure levels from those cohort subjects who fully participated. While the duration of breast-feeding was associated with the child’s serum-PFAS concentrations [14], this parameter was unrelated to the antibody concentrations. No important confounders were identified among a wide range of social and demographic parameters, and adjustments therefore included only sex and age and, for the age-7 data, the type of booster vaccination at age 5. These covariates affected the results to a negligible degree only.

As can be seen from the (previously published) left-hand columns of Table 1, the log model yielded lower BMDs and BMDLs, and the model-dependence was similar for tetanus and diphtheria antibody concentrations as outcome variables. When using the linear slope and a BMR of 5%, the BMDL was about 1.3 ng/mL and 0.3 ng/mL for PFOS and PFOA, respectively. The piecewise linear curve was steeper at low exposures, it showed BMDL results about half of those obtained by the linear dose-response curve, while the logarithmic curve showed even lower results. In addition, results were higher at a BMR of 10%. Mutual adjustment (right-hand columns of Table 1) resulted in higher BMD values, while changes in BMDL were rather small and much lesser than the differences seen between the dose-response models. All dose-response models had normally distributed residuals with a homogeneous scatter. Table 2 shows that the various models almost equally well fit the data. The piecewise linear generally tended to show slightly better fit values, but none of the differences observed reached statistical significance.

**Table 1.**
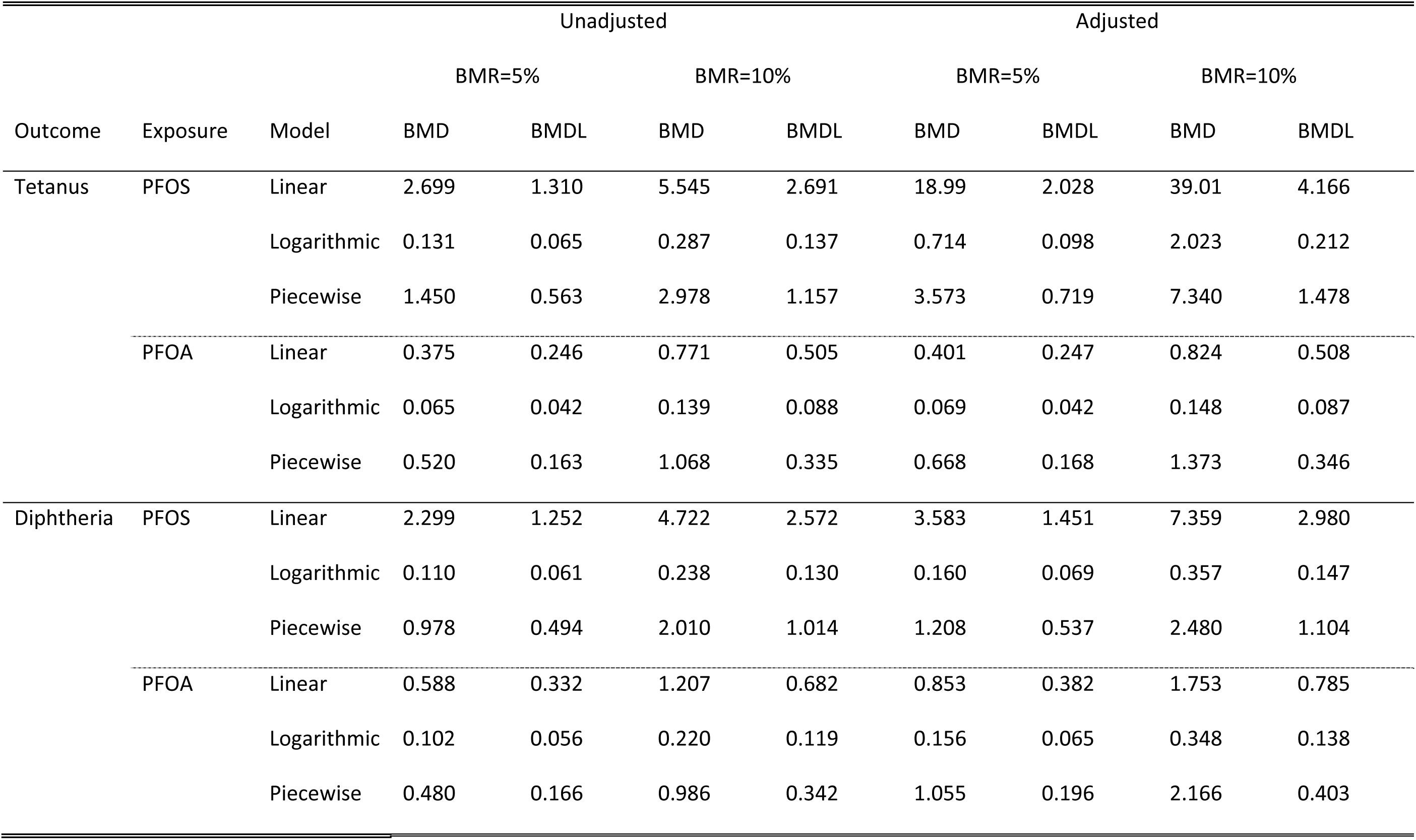
Benchmark results for the age-5 serum concentrations of PFOS and PFOA in regard to tetanus and diphtheria antibody concentrations at age 7 years (after four vaccinations). Unadjusted benchmark results are followed by results mutually adjusted for the two PFAS concentrations.

**Table 2.** Fit of dose-response models used in Table 1. The fit was estimated as minus two times the log-maximum likelihood function (–2 ln (L)), where a smaller value indicates a better fit. The fit is given for the whole dose range and at low dose and before and after adjustment for the other PFAS.

| Outcome | Exposure | Model | Unadjusted |  | Adjusted |  |
| --- | --- | --- | --- | --- | --- | --- |
|  |  |  | Full Range | Low dose | Full range | Low dose |
| Tetanus | PFOS | Linear | 1719.81 | 313.78 | 1713.18 | 311.11 |
|  |  | Log | 1719.30 | 313.75 | 1713.10 | 311.02 |
|  |  | Piecewise | 1719.54 | 313.54 | 1713.04 | 310.99 |
|  | PFOA | Linear | 1712.43 | 391.53 | 1712.24 | 391.03 |
|  |  | Log | 1712.88 | 391.63 | 1712.77 | 391.25 |
|  |  | Piecewise | 1712.33 | 391.64 | 1712.09 | 391.07 |
| Diphtheria | PFOS | Linear | 1656.86 | 314.00 | 1655.01 | 313.71 |
|  |  | Log | 1655.96 | 313.38 | 1654.44 | 313.21 |
|  |  | Piecewise | 1655.77 | 313.10 | 1654.08 | 312.92 |
|  | PFOA | Linear | 1656.15 | 362.37 | 1654.12 | 361.39 |
|  |  | Log | 1656.14 | 362.39 | 1654.31 | 361.48 |
|  |  | Piecewise | 1656.12 | 362.30 | 1654.11 | 361.41 |

Using the prenatal exposure levels and the age-5 antibody outcomes, somewhat higher results were obtained (Table 3). The differences between the models and the effect of mutual adjustment followed the same pattern as the postnatal data (Table 1), although the differences tended to be smaller for this data set. Table 4 shows the calculated fits for the different models for this data set. The strongest difference in fit was seen for the effect of prenatal PFOA exposure on diphtheria-specific antibody concentrations at age 5. Here the pricewise linear model had superior fit both in the whole dose-range and at low doses and for this association the piecewise model produced benchmark results that were comparable to those obtained by the logarithmic model.

**Table 3.** Benchmark results for the prenatal PFAS concentrations in regard to antibody concentrations at age 5 years (pre-booster). Unadjusted Benchmark results are followed by results mutually adjusted for the two PFAS concentrations.

| Outcome | Exposure | Model | Unadjusted |  |  |  | Adjusted |  |  |  |
| --- | --- | --- | --- | --- | --- | --- | --- | --- | --- | --- |
|  |  |  | BMR=5% |  | BMR=10% |  | BMR=5% |  | BMR=10% |  |
|  |  |  | BMD | BMDL | BMD | BMDL | BMD | BMDL | BMD | BMDL |
| Tetanus | PFOS | Linear | 7.247 | 2.858 | 14.89 | 5.870 | ∞ | 4.758 | ∞ | 9.773 |
| | | Logarithmic | 0.338 | 0.135 | 0.818 | 0.298 | ∞ | 0.246 | ∞ | 0.570 |
| | | Piecewise | 2.623 | 1.062 | 5.388 | 2.182 | ∞ | 1.662 | ∞ | 3.415 |
|  | PFOA | Linear | 0.550 | 0.316 | 1.130 | 0.649 | 0.589 | 0.310 | 1.210 | 0.637 |
|  |  | Logarithmic | 0.136 | 0.080 | 0.299 | 0.172 | 0.134 | 0.075 | 0.294 | 0.159 |
|  |  | Piecewise | 0.248 | 0.134 | 0.510 | 0.275 | 0.246 | 0.127 | 0.506 | 0.261 |
| Diphtheria | PFOS | Linear | 2.386 | 1.559 | 4.900 | 3.201 | 3.071 | 1.735 | 6.309 | 3.564 |
|  |  | Logarithmic | 0.120 | 0.077 | 0.262 | 0.164 | 0.158 | 0.084 | 0.352 | 0.181 |
|  |  | Piecewise | 1.617 | 0.829 | 3.321 | 1.703 | 2.486 | 0.943 | 5.107 | 1.938 |
|  | PFOA | Linear | 0.654 | 0.339 | 1.342 | 0.696 | 2.595 | 0.504 | 5.330 | 1.034 |
|  |  | Logarithmic | 0.139 | 0.080 | 0.305 | 0.171 | 0.332 | 0.108 | 0.801 | 0.235 |
|  |  | Piecewise | 0.150 | 0.097 | 0.307 | 0.199 | 0.213 | 0.115 | 0.437 | 0.236 |

**Table 4.** Fit of dose response models used in Table 3. The fit was estimated as minus two times the log-maximum likelihood function (−2 ln (L)), where a smaller value indicates a better fit. The fit is given for the whole dose range and at low dose and before and after adjustment for the other PFAS.

| Outcome | Exposure | Model | Unadjusted |  | Adjusted |  |
| --- | --- | --- | --- | --- | --- | --- |
|  |  |  | Full Range | Low dose | Full range | Low dose |
| Tetanus | PFOS | Linear | 3481.00 | 859.75 | 3474.77 | 856.25 |
|  |  | Log | 3480.53 | 859.20 | 3474.76 | 856.30 |
|  |  | Piecewise | 3480.40 | 859.29 | 3474.77 | 856.25 |
|  | PFOA | Linear | 3477.21 | 822.35 | 3477.13 | 822.14 |
|  |  | Log | 3475.61 | 824.75 | 3475.61 | 824.78 |
|  |  | Piecewise | 3475.88 | 820.94 | 3475.88 | 820.96 |
| Diphtheria | PFOS | Linear | 3566.54 | 831.18 | 3564.76 | 828.12 |
|  |  | Log | 3566.48 | 830.89 | 3565.26 | 828.41 |
|  |  | Piecewise | 3566.17 | 831.03 | 3564.71 | 828.23 |
|  | PFOA | Linear | 3573.02 | 838.51 | 3566.61 | 836.47 |
|  |  | Log | 3570.50 | 840.06 | 3565.92 | 838.93 |
|  |  | Piecewise | 3566.53 | 835.98 | 3562.84 | 835.16 |

Results of the sensitivity analyses are shown in Table 5. Assuming a low-dose threshold in each of the dose-response models, the differences between them are minimized, as might be expected when a flat slope is inserted in the low dose range, thereby minimizing the differences between the curves in the low dose range. Again, mutual adjustment had a very minor effect on the BMDL results.

**Table 5:** Benchmark results for PFAS concentration for antibody concentrations for the two sets of data using conservative models with no-effect below lowest observed exposure levels. Unadjusted benchmark results (examples) are followed by results adjusted for the other PFAS concentration.

|  |  |  | Unadjusted |  |  |  | Adjusted |  |  |  |
| --- | --- | --- | --- | --- | --- | --- | --- | --- | --- | --- |
|  |  |  | BMR=5% |  | BMR=10% |  | BMR=5% |  | BMR=10% |  |
| Postnatal | Exposure | model | BMD | BMDL | BMD | BMDL | BMD | BMDL | BMD | BMDL |
| Tetanus | PFOA | Linear | 1.702 | 1.573 | 2.098 | 1.832 | 1.728 | 1.574 | 2.151 | 1.835 |
|  |  | Log | 1.479 | 1.425 | 1.650 | 1.532 | 1.489 | 1.424 | 1.671 | 1.530 |
|  |  | Piecewise | 1.874 | 1.495 | 2.450 | 1.672 | 2.044 | 1.501 | 2.800 | 1.684 |
| Diphtheria | PFOS | Linear | 8.476 | 7.429 | 10.90 | 8.749 | 9.760 | 7.628 | 13.54 | 9.157 |
|  |  | Log | 6.964 | 6.616 | 7.887 | 7.107 | 7.327 | 6.674 | 8.740 | 7.235 |
|  |  | Piecewise | 7.191 | 6.700 | 8.259 | 7.251 | 7.437 | 6.748 | 8.764 | 7.349 |
| Prenatal |  |  |  |  |  |  |  |  |  |  |
| Tetanus | PFOA | linear | 0.929 | 0.695 | 1.509 | 1.028 | 0.968 | 0.689 | 1.589 | 1.016 |
|  |  | log | 0.566 | 0.490 | 0.791 | 0.617 | 0.563 | 0.482 | 0.785 | 0.598 |
|  |  | broken | 0.630 | 0.514 | 0.895 | 0.656 | 0.628 | 0.507 | 0.891 | 0.643 |
| Diphtheria | PFOS | linear | 4.274 | 3.447 | 6.788 | 5.089 | 4.959 | 3.623 | 8.197 | 5.452 |
|  |  | log | 2.234 | 2.110 | 2.645 | 2.363 | 2.345 | 2.131 | 2.904 | 2.410 |
|  |  | broken | 3.446 | 2.709 | 5.088 | 3.574 | 4.195 | 2.814 | 6.627 | 3.790 |

## Discussion

Benchmark calculations are thought to provide an approximate threshold level which can be interpreted as a parallel to the No-Observed Adverse Effect level (NOAEL) from experimental studies [34]. The lower confidence limit(the BMDL) takes into account the uncertainty in the estimation of the relation between dose and effect for a given dose-response model [34]. The present report complements our previous report on BMD results for postnatal PFAS exposure [30] by adding calculations for prenatal PFAS exposures and for mutual adjustments for PFOS and PFOA exposures. Regulatory agencies have so far only used benchmark dose results obtained from animal studies of PFAS toxicity [26, 36-38], but our new results provide more extensive data to allow comparison between laboratory results and epidemiological findings.

The size and homogeneity of the study population and the high participation rate are major strengths [16], as is the fact that neighbourhood or occupational industrial exposures and water contamination do not affect the exposure profile in this community. PFOS and PFOA were the predominant PFASs in this population, where marine food contamination is an important source. As the interest mainly focuses on the two major PFASs, we present simple regression models, although our structural equation model analyses suggest that the overall effects of PFASs on the antibodies were stronger than most individual effects [16]. Thus, imprecision in the individual exposure measurement for the PFASs could result in an underestimation of their toxicity [39].

We chose to focus on the specific IgG antibody concentrations against two toxoid vaccines as well defined immune system responses. The pre-booster concentration at age 5 represents the long-term response after the first three vaccinations against tetanus and diphtheria, while the level two years later takes into account the response following the age-5 booster. Other clinical outcomes that reflect immune functions may be less sensitive and also more difficult to assess in a standardized way. For example, hospitalization of 363 children for infectious disease (such as middle ear infection, pneumonia, and appendicitis) up to an average age of about 8 years was not associated with PFOS and PFOA concentrations in serum from pregnant women from the Danish National Birth Cohort [40]. While multiple social, demographic, and other factors may have affected these results, hospitalization does not seem to be a sensitive or appropriate test in detecting immune system dysfunction, at least to the extent that it reflects an impact of prenatal PFAS exposure. In contrast, more recent studies have documented increased frequencies of common infections in small children at elevated background levels of PFASs in maternal pregnancy serum [41, 42]. In support of these observations, white blood cells from human volunteers showed effects at PFOS concentrations in the medium of 0.1 μg/mL)100 ng/mL), which was the lowest concentration tested [29]. This concentration is similar to the highest serum levels seen at background exposures [9].

While vaccine responses depend on the adaptive immune system, which undergoes important development during infancy, the most vulnerable time window for PFAS immunotoxicity is unknown. However, in a recent study, we modeled serum concentrations during infancy and found that levels at ages 3 and 6 months appeared to be at least as strong predictors of decreased vaccine responses at age 5 years as was the maternal concentrations [17]. The time dependence is difficult to explore in detail due to the complexity of obtaining blood samples from small children. In addition, the long elimination half-life of both PFOS and PFOA [43] means that the impact of PFAS transfer through human milk will still be apparent in serum concentrations at age 5 [14]. Thus, prenatal and early postnatal exposures will remain in the body for several years, and any age-dependent effects are difficult to separate. However, the availability of serum concentrations only at two points in time likely results in some underestimation of the associations at the most sensitive age.

An important weakness of epidemiological studies is the mixed exposures. Among PFASs studied so far, PFOS and PFOA generally occur in serum in the largest concentrations [9], and their immunotoxic effects are well documented [7], although their adverse outcome pathways may differ [29]. In our past studies, we have utilized structural equation models to assess the total impact of the (mixed) PFAS exposures, in part to take into account the imprecision of the exposure measurements for individual PFASs [16, 28]. Joint effects of the major PFASs seemed stronger than those that could be ascribed to single compounds, and more than one PFAS therefore may contribute to the lowering of antibody responses. Given the strong experimental support for immunotoxicity of both PFOS and PFOA [7], the BMD analyses for each of the two would seem appropriate, as would mutual adjustments, as presented in the present study. However, the current regression models did not allow for a possible interaction effect between the two PFAS concentrations. If such effects are present, the calculated Benchmark results may not be as protective and intended.

Given that the BMDL is calculated as the statistical 95% lower bound of the BMD, it depends on the statistical uncertainty of the BMD determination. When covariates are added to the equation, greater uncertainty will often occur, thus resulting in a greater BMD and a lower BMDL (relative to the BMD). In the current application, we adjusted the BMD results for the effect of a covariate strongly correlated with the exposure of interest. General statistical advice is to avoid inclusion of strongly correlated covariates, as the results may be over-adjusted and unstable. If inference is based on the *p* value, harmful exposures may be overlooked or disregarded as a result of inappropriate adjustment. However, these concerns may not extend to BMD analyses. Here the inference is based on the lower confidence limit, the BMDL, and over-adjustment with increased variance will likely lead to lower BMDL results that may cancel out the BMD increase. The lowered BMDL can be considered in accordance with the precautionary principle [31] and may argue against current wisdom to avoid adjustment for closely related co-exposures when BMD calculations are the purpose.

As demonstrated by our results, the choice of dose-response model results in differing BMD results, especially in epidemiological studies where unexposed controls are usually missing [31]. In the absence of prior knowledge regarding the most relevant curve function, we used two common curve shapes (linear and logarithmic) to explore the dependence on the two assumptions. The curves fit the data about equally well. As a result, no statistical justification is available for choosing one set of results above the others. The linear curve is often used as a default, and we therefore also examined a model with a piecewise linear shape that allowed a different slope below the median exposure level. For each of the two PFASs, these sensitivity analyses showed that the BMDL results remained low. As also anticipated, the 5% BMR resulted in BMDL values somewhat below those for 10%, but differences between the curve shapes were of similar magnitude also at the higher BMR value. Again, these tendencies were replicated in the mutual adjustment results.

As a consequence of the relatively steep dose-response relationships, the BMD results were often lower than the lowest observed exposure level. Consequently, some results depended on a part of the dose-response curve for which the data do not hold any information. The dose-response model that allowed a different slope at exposures below the median does not resolve this concern. In our previous study [30], we also included sensitivity analyses, where we assumed that no change in the antibody occurred below the lowest observed exposure level. As expected, these models yielded higher BMD results, but the increase in BMDL was not substantial. Interpretation of these results must be cautious due to the questionable plausibility of the low-dose flat curve shape.

Given the differences between BMDL values for different dose-response models, different adjustments and different ages at exposure and outcomes, the present study identifies a range of BMDLs that would be appropriate for estimation of exposure limits. An approximate BMDL of 1 ng/mL serum for both PFOS and PFOA (somewhat higher for PFOS and lower for PFOA) would seem to appropriate. As the BMDL assumes equal sensitivity within the population studied, current guidelines [20, 21] require that the BMDL be divided by an uncertainty factor of 10 to take into account the existence of subjects with increased vulnerability. A reference concentration of about 0.1 ng/mL could then be used as the serum-based target. This concentration is below most human serum concentrations of PFOS and PFOA reported [9], and it is also greatly below previously derived BMDLs from animal toxicity tests [26]. Still, recent data on mammary gland development suggests this outcome as an additional highly sensitive target in rodents [44, 45]. When applying a standard uncertainty factor of 100, the resulting reference level is quite similar to the one that we have calculated for immunotoxicity in humans. Future refinement of exposure limits for PFASs should take into account these insights into BMDL calculations.

## Acknowledgments

This study was supported by the National Institute of Environmental Health Sciences, NIH (R01ES012199, R01ES026596, and P42ES027706); and the Danish Environmental Protection Agency as part of the environmental support program DANCEA (Danish Cooperation for Environment in the Arctic) and the Dynamical Systems Interdisciplinary Network, University of Copenhagen. We thank our close colleagues, Pal Weihe and Carsten Heilmann for their support during this effort as a contribution to our overall collaborative study.

